# The major role of *sarA* in limiting *Staphylococcus aureus* extracellular protease production is correlated with decreased virulence in diverse clinical isolates in osteomyelitis

**DOI:** 10.1101/2022.11.07.515558

**Authors:** Mara J. Campbell, Karen E. Beenken, Aura M. Ramirez, Mark S. Smeltzer

## Abstract

We previously demonstrated that MgrA, SarA, SarR, SarS, SarZ, and Rot bind at least three of the four promoters associated with genes encoding primary extracellular proteases in *Staphylococcus aureus*. We also showed that mutation of *sarA* results in a greater increase in protease production, and decrease in biofilm formation, than mutation of the loci encoding any of these other proteins. However, these conclusions were based on *in vitro* studies. Thus, the goal of the experiments reported here was to determine the relative impact of the regulatory loci encoding these proteins *in vivo*. To this end, we compared the virulence of *mgrA, sarA, sarR, sarS, sarZ*, and *rot* mutants in a murine osteomyelitis model. Mutants were generated in the methicillin-resistant USA300 strain LAC and the methicillin-sensitive USA200 strain UAMS-1. As assessed based on an overall osteomyelitis pathology score derived from the incidence of bone fracture, bacterial burdens in the bone, cortical bone destruction, and reactive bone formation, mutation of *mgrA* and *rot* limited virulence to a statistically significant extent in UAMS-1, but not in LAC. In contrast, the *sarA* mutant exhibited reduced virulence in both strains. This illustrates the importance of considering diverse clinical isolates when evaluating the impact of regulatory mutations on virulence. The reduced virulence of the *sarA* mutant was correlated with reduced cytotoxicity for osteoblasts and osteoclasts, reduced biofilm formation, and reduced sensitivity to the antimicrobial peptide indolicidin, all of which were directly attributable to increased protease production in both LAC and UAMS-1. This suggests that these *in vitro* phenotypes, either alone or in combination with each other, may be useful in prioritizing additional mutants for *in vivo* evaluation. Most importantly, they illustrate the significance of limiting protease production *in vivo* in S. *aureus*, and confirm that SarA plays the primary role in this regard.

**Author Summary:** *Staphylococcus aureus* causes a diverse array of infections due to its ability to produce an arsenal of virulence factors. Among these are extracellular proteases, which serve several purposes on behalf of the bacterium. However, it has become increasingly apparent that it is also critical to limit the production of these proteases to prevent them from compromising the *S. aureus* virulence factor repertoire. Many regulatory loci have been implicated in this respect, but it is difficult to draw relative conclusions because few reports have made direct comparisons, and fewer still have done so *in vivo*. We addressed this by assessing the impact on virulence of six regulatory loci previously implicated in protease production. We did this in the clinical context of osteomyelitis using mutants generated in two divergent clinical isolates. Our results confirm significant strain-dependent differences, reinforcing the importance of considering such diverse clinical isolates when evaluating targets for potential therapeutic intervention. In this respect, only mutation of *sarA* attenuated virulence in both strains. This illustrates the importance of limiting protease production as a means of post-translational regulatory control in *S. aureus* and confirms that *sarA* plays a predominant role in this regard.

## Introduction

Although many bacterial pathogens have been etiologically associated with osteomyelitis, *Staphylococcus aureus* is overwhelmingly the most common cause and the pathogen that causes the most damage to the bone and surrounding tissues [1–3]. Osteomyelitis is a uniquely problematic form of *S. aureus* infection owing to its complex pathology and intrinsic resistance to conventional antibiotic therapy [4,5]. One reason for this intrinsic resistance is that most infections are not diagnosed until they have progressed to a chronic stage in which the bone and vasculature at the site of infection has been compromised, thus limiting systemic antibiotic delivery [2,6]. Moreover, a key characteristic of osteomyelitis is formation of a biofilm, which confers a therapeutically relevant level of intrinsic resistance to all antibiotics at both an aggregate and cellular level [7–9]. *S. aureus* can also invade, survive, and even replicate inside osteoblasts and osteoclasts, thus providing further protection from antibiotics and host defenses [10–13]. As a result, the effective treatment of osteomyelitis requires a multidisciplinary approach that includes long-term systemic antibiotic therapy and surgical debridement, often accompanied by some form of local, matrix-based antibiotic delivery directly to the site of infection [5,6,14,15]. Even after such intensive medical and surgical intervention, the recurrence rate in highly complex cases can be as high as 20-30%, often resulting in amputation [4,6,8,16].

Mutation of the staphylococcal accessory regulator (*sarA*) limits biofilm formation to a degree that can be directly correlated with increased antibiotic susceptibility [17–20] and limits virulence in a murine osteomyelitis model [21–24]. This attenuation is attributable in large part to the increased production of extracellular proteases in *sarA* mutants and the corresponding decrease in the abundance of multiple *S. aureus* virulence factors [23–27]. These results led to the hypothesis that, while extracellular proteases contribute to virulence by promoting tissue invasion, the acquisition of nutrients, and avoidance of host defenses [28,29], it is equally important that their production be limited such that they serve these purposes on behalf of the bacterium without compromising the virulence factor repertoire of *S. aureus* [21,23–26,30].

The importance of controlling the production of extracellular proteases is reflected in the number of *S. aureus* regulatory loci that have been shown to modulate their production to a degree that impacts potentially important osteomyelitis-related phenotypes including biofilm formation [23–25,31–39]. Indeed, Fey *et al* [40] used casein agar plates to screen the Nebraska Transposon Mutant Library (NTML) and identified 62 mutants that exhibited differences in protease activity, with 12 exhibiting increased activity. Similarly, based on transcriptional analysis Gimza et al [37] identified more than 50 putative *S. aureus* regulatory elements that impact transcription of one or more of the four genes or operons encoding the proteases aureolysin, SspA/SspB, ScpA, or SplA-F.

In a previous report, we used an DNA-based capture assay to identify regulatory proteins that bind the promoter regions associated with each of the four genes/operons encoding these proteases. The results of these studies confirmed that MgrA, SarA, SarS, and Rot bind to *cis* elements associated with all four promoters, while SarR and SarZ bind three of four protease-associated DNA baits, the exception in both cases being the promoter associated with *scpA* [23]. SarA was the most abundant protein captured by all four baits, and mutation of *sarA* resulted in a significantly greater increase in overall protease production than mutation of any of the regulatory loci encoding these other proteins [23]. It also resulted in a more significant reduction in biofilm formation and the accumulation of high-molecular weight proteins. The phenotypic impact of mutating *sarA* on protease production was also evident irrespective of the functional status of any of these other regulatory loci [23].

These results led us to conclude that *sarA* plays the predominant role in ensuring that the production of extracellular proteases does not exceed levels that compromise the *S. aureus* virulence factor repertoire [23,25]. However, this conclusion was based solely on the results of *in vitro* studies, and no *in vitro* condition can be assumed to reflect *in vivo* relevance. This is particularly true since *S. aureus* is a unique bacterial pathogen with the ability to cause infection in diverse microenvironments within the host [41]. For instance, bone is intrinsically hypoxic, and oxygen availability is known to impact regulatory circuits and the production of multiple virulence factors in *S. aureus* [42]. To our knowledge, *sarA* is the only one of these regulatory loci that has been examined in the pathogenesis of osteomyelitis. Thus, the relative role of these other regulatory loci *in vivo* in this important clinical context remains to be determined.

To address this, we generated mutations in *mgrA, sarA, sarS, sarR, sarZ* and *rot* and evaluated the impact on virulence in a murine osteomyelitis model. These experiments were done with mutants generated in the methicillin-resistant USA300 strain LAC and the methicillin-sensitive USA200 strain UAMS-1 to account for the genotypic and phenotypic diversity among *S. aureus* clinical isolates. The results of *in vivo* comparisons confirmed that mutation of *sarA* limited the virulence of both strains and that this was not true of any other regulatory mutant. This finding is consistent with the hypothesis that limiting protease production plays an important role in post-translational regulation in S. *aureus*, and that SarA plays a predominant role in this regard. This suggests that *sarA* or its protein product could be a viable target for the development of alternative methods that could potentially be used to limit the pathology and overcome the therapeutic recalcitrance of *S. aureus* osteomyelitis.

## Results

### Impact of regulatory loci on virulence in LAC

The relative virulence of *mgrA*, *sarA*, *sarR*, *sarS*, *sarZ* and *rot* mutants was assessed in the USA300 strain LAC in two independent experiments. The only two groups in which no fractures were observed in any mice in either experiment were those infected with the *sarA* and *rot* mutants (Fig 1A). Quantitative μCT analysis of intact bones for cortical bone destruction (Fig 1B) and reactive new bone (callous) formation (Fig 1C) demonstrated that the only statistically significant reduction by comparison to mice infected with LAC was in mice infected with the *sarA* mutant. Mice infected with the *sarA* mutant were also the only group with a significantly reduced bacterial burden (Fig 1D). When the incidence of fractured bones was included to derive an overall osteomyelitis score (OM) for all animals, the only statistically significant difference was in the group infected with the *sarA* mutant (Fig 1E).

**Fig 1.**
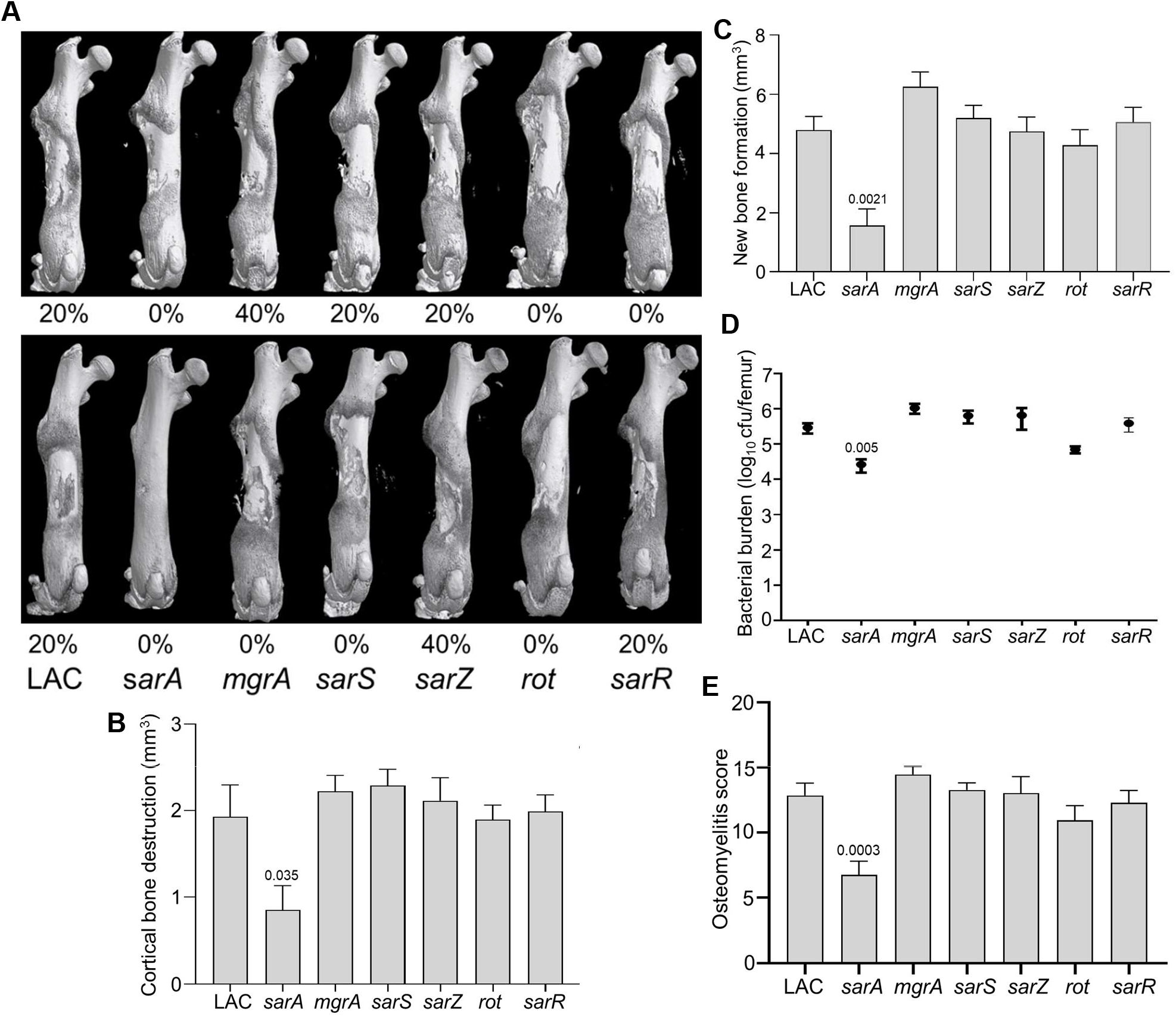
Mutation of *sarA* in LAC attenuates virulence in a murine osteomyelitis model to a greater extent than mutation of any other regulatory locus. A) Femurs were harvested from mice 14 days post-infection and imaged by μCT. Top and bottom panels are randomly chosen images mice from each experimental group obtained in two independent experiments. Numbers below each panel denote the percentage of broken bones observed in each experimental group. B and C) Quantitative analysis of cortical bone destruction (B) and new bone formation (C) of intact bones. D) Bacterial burdens were determined by homogenization of fractured and unfractured bones and serial dilution of the homogenates prior to plating on TSA. Results are reported as the average ± the standard error of the mean (SEM). E) Overall osteomyelitis (OM) scores were calculated to allow consideration of intact and broken bones. In all cases, statistical significance was assessed by one-way ANOVA. Numbers indicate p-values by comparison to the results observed with mice infected with the LAC parent strain.

### Impact of regulatory loci on osteomyelitis-associated phenotypes in LAC

The results observed with LAC and its regulatory mutants in our osteomyelitis model were directly reflected in studies assessing the impact of regulatory mutants on osteoblast and osteoclast cytotoxicity. Specifically, conditioned medium (CM) from LAC was cytotoxic for both cell types, and the only mutation that significantly reduced this cytotoxicity was *sarA* (Fig 2A and B). Additionally, cytotoxicity was fully restored in the LAC *sarA* mutant by eliminating the production of extracellular proteases (Fig 2A and B). Mutation of *sarA* also limited biofilm formation in LAC to a statistically significant degree, and this was also reversed by eliminating the ability of the *sarA* mutant to produce extracellular proteases (Fig 2C). Mutation of *rot* also reduced the capacity of LAC to form a biofilm, and this effect was also reversed by eliminating the ability of the *rot* mutant to produce extracellular proteases (Fig 2C). Growth in the presence of indolicidin, which we used as a representative antimicrobial peptide (AMP), was significantly increased in the *sarA* mutant in a protease dependent manner (Fig 2D). No other regulatory mutants had a significant increase in growth in the presence of indolicidin by comparison to LAC, but mutation of *sarZ* resulted in a significant reduction in growth (Fig 2D). Survival in whole human blood was increased to a significant extent in LAC *sarA* mutant (Fig 2E). However, eliminating the production of extracellular proteases in the *sarA* mutant resulted in only a slight reduction in growth in whole human blood that was not statistically significant when compared to the *sarA* mutant (Fig 2E).

**Fig 2.**
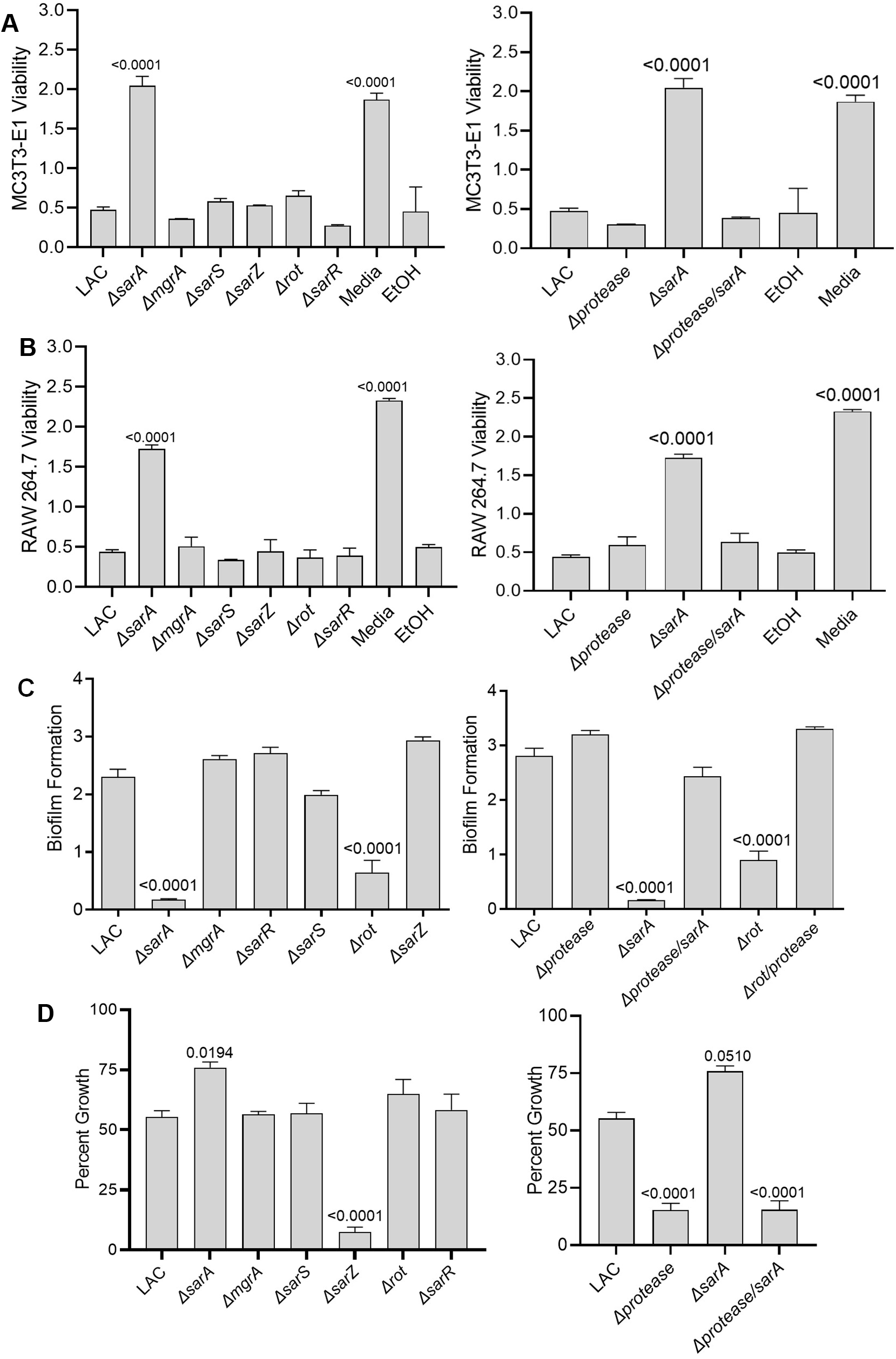
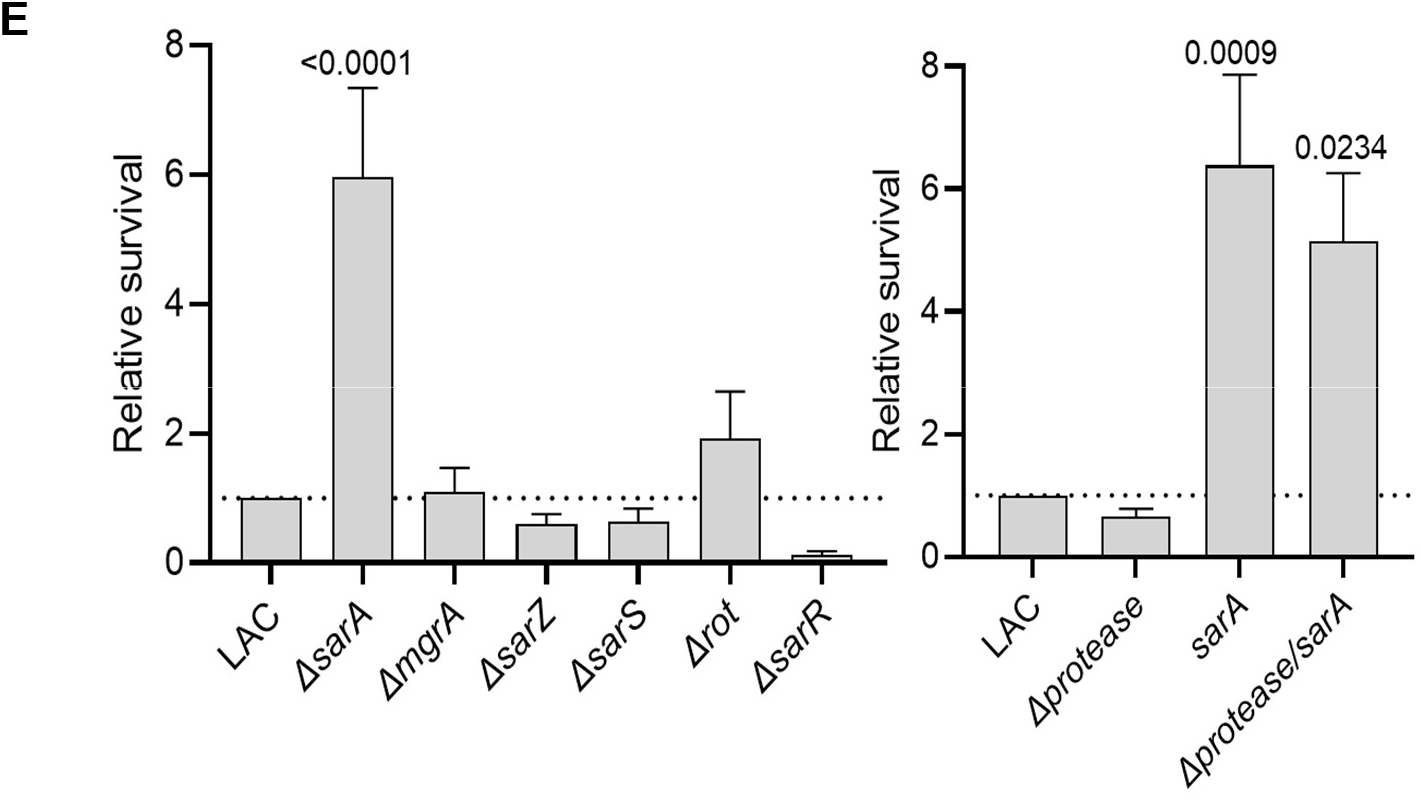
Mutation of *sarA* in LAC has the greatest effect on osteomyelitis related phenotypes owing primarily to its impact on protease production. A and B) Cytotoxicity of conditioned medium (CM) from overnight cultures of LAC and isogenic mutants was assessed using MC3T3-E1 (A) and RAW264.7 cells (B) as surrogates for osteoblasts and osteoclasts, respectively. CM, sterile bacterial culture media (negative control), or ethanol (positive control) was mixed in a 1:1 ratio with the appropriate cell culture medium for cytotoxicity assays. C) Biofilm formation was assessed using a microtiter plate assay. The impact of mutating *rot* on biofilm formation was statistically different from that of mutating *sarA* (p = 0.0214). D) For each strain, 1×10^6^ cfu was inoculated into TSB with or without 10 μg/mL indolicidin and incubated overnight with shaking. Growth was measured by OD_600_ relative to a DMSO control for each mutant. E) 1×10^5^ cfu of bacterial cells from exponential phase cultures were mixed with 1.0 ml of whole human blood. A sample was taken immediately after mixing and after a 3 hr incubation. Percent survival of each mutant was calculated and standardized relative to the results observed with LAC. The difference between the results observed with the *sarA* mutant and its protease-deficient derivative were not statistically significant. In all cases, statistical significance was assessed by one-way ANOVA. Numbers indicate p-values by comparison to the results observed with the LAC parent strain.

### Impact of regulatory loci on virulence in UAMS-1

Mutation of *sarA* also attenuated the virulence of UAMS-1, but the results were not as definitive as those observed in LAC. For instance, no broken bones were observed in either of two independent experiments in mice infected with the *sarA, mgrA*, or *sarR* mutants (Fig 3A). Moreover, while clear downward trends were observed with UAMS-1 *sarA* and *mgrA* mutants, no statistically significant differences were observed in cortical bone destruction (Fig 3B). Mutation of *sarA, mgrA* or *rot* did result in a statistically significant reduction in new bone formation (Fig 3C), and mutation of *mgrA* or *rot* resulted in significantly decreased bacterial burdens while mutation of *sarA* resulted in a downwards trend that did not reach statistical significance (p-value 0.0526) (Fig 3D). When all of the *in vivo* data was combined to generate an overall osteomyelitis score, mutation of *sarA, mgrA*, or *rot* were all found to result in a statistically significant reduction in virulence (Fig 3E).

**Fig 3.**
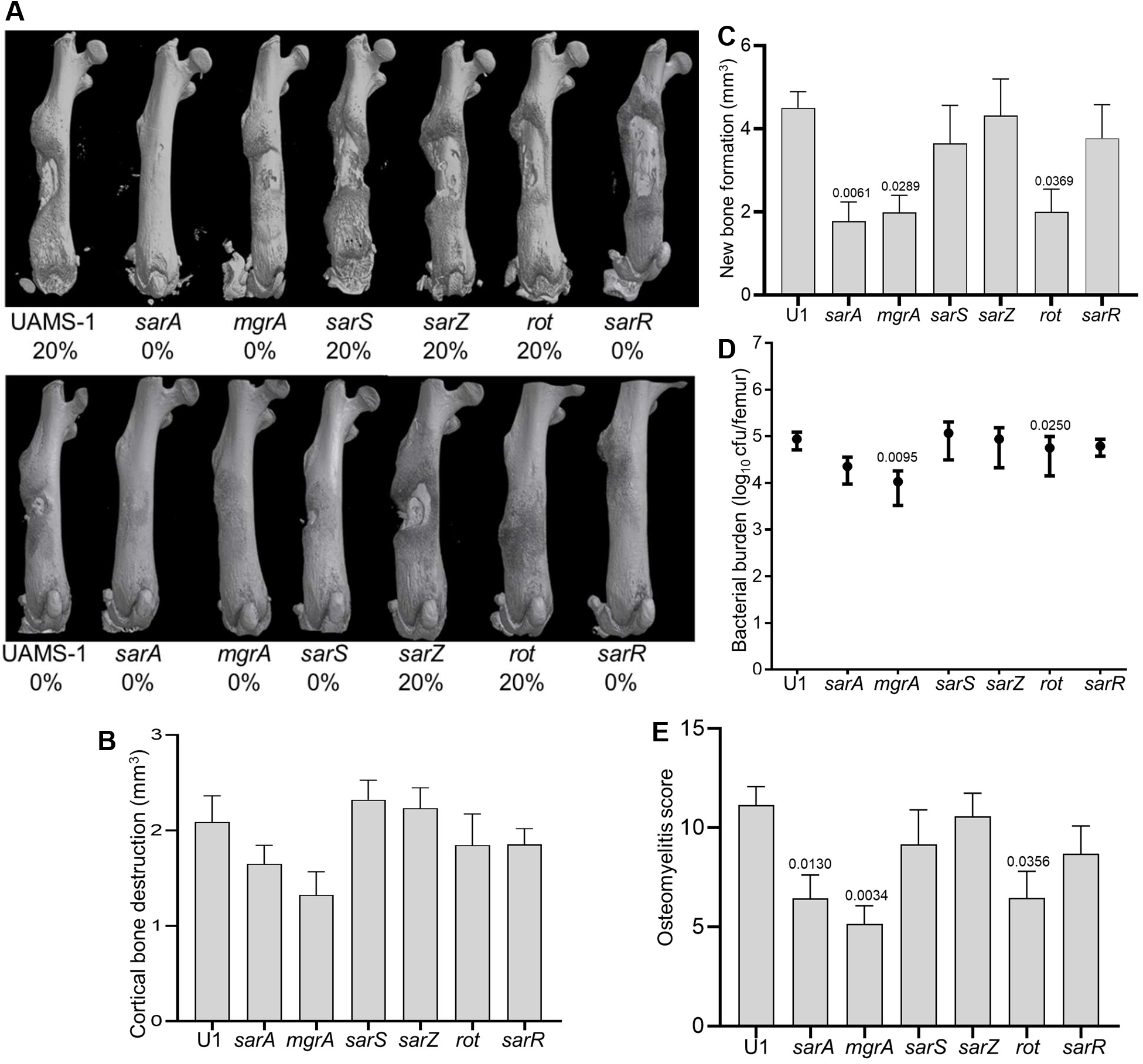
Mutation of *sarA, mgrA*, or *rot* in UAMS-1 results in attenuation in a murine osteomyelitis model. A) Femurs were harvested from mice 14 days post-infection and imaged by μCT. Top and bottom panels are randomly chosen images mice from each experimental group obtained from two independent experiments. Numbers below each panel denote the percentage of broken bones observed in each experimental group. B and C) Quantitative analysis of cortical bone destruction (B) and new bone formation (C) of intact bones. D) Bacterial burdens were determined by homogenization and plating after μCT and reported as the average ± the standard error of the mean (SEM). No statistically significant difference was observed in bacterial burdens with the *sarA* mutant, but a downward trend was evident (p = 0.0526). E) Overall osteomyelitis (OM) scores were calculated to allow inclusion of both intact and broken bones. In all cases, statistical significance was assessed by one-way ANOVA. Numbers indicate p-values by comparison to the results observed with mice infected with the UAMS-1 parent strain.

### Impact of regulatory loci on osteomyelitis-associated phenotypes in UAMS-1

As was observed in LAC, only mutation of *sarA* resulted in a significant increase in cytotoxicity for osteoblasts and osteoclasts (Fig 4A and B), and this was directly correlated with the increased production of extracellular proteases (Fig 4A and B). Similarly, mutation of *sarA* resulted in decreased biofilm formation (Fig 4C) and increased growth in the presence of indolicidin (Fig 4D), both of which were defined by the increased production of extracellular proteases (Fig 4C and D). As in LAC, survival in whole human blood was also increased in a UAMS-1 *sarA* mutant, and this phenotype was not protease dependent (Fig. 4E). In contrast to LAC, mutation of *rot* did not result in a statistically significant decrease in biofilm formation (Fig 4C) and mutation of *sarZ* did not result in significantly decreased growth in the presence of indolicidin (Fig 4D).

**Fig 4.**
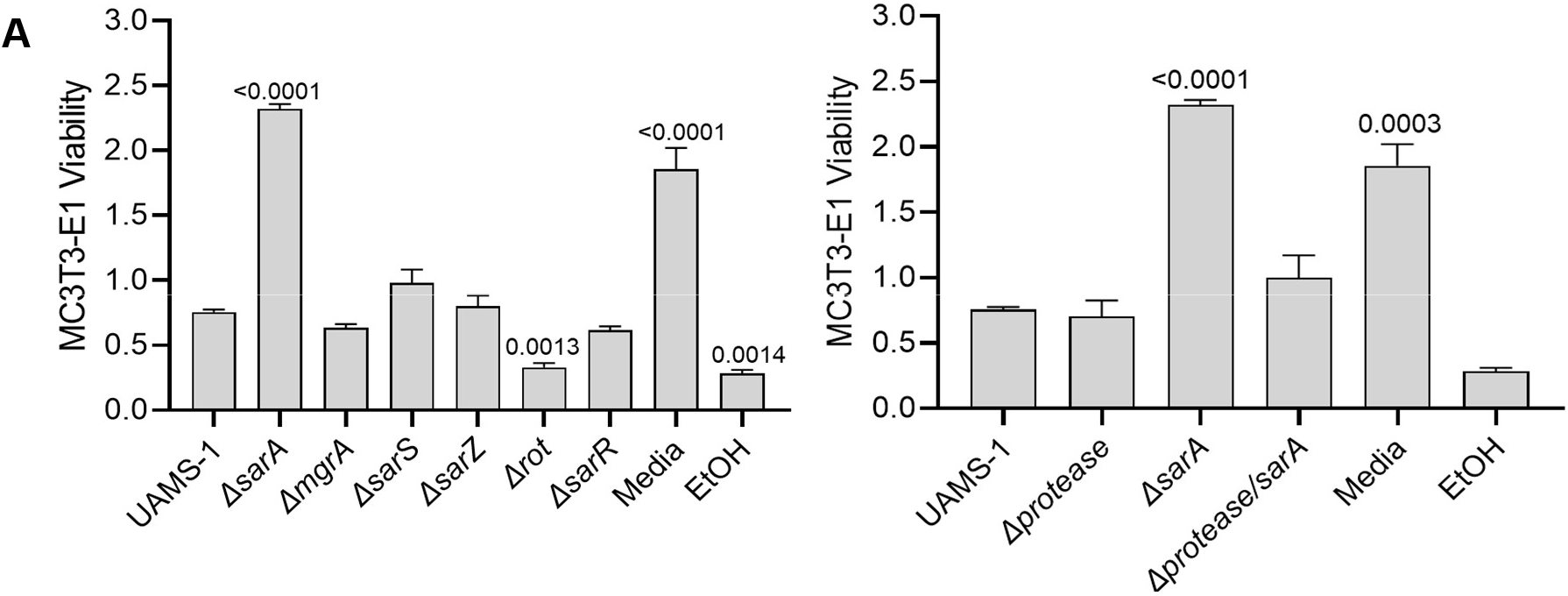
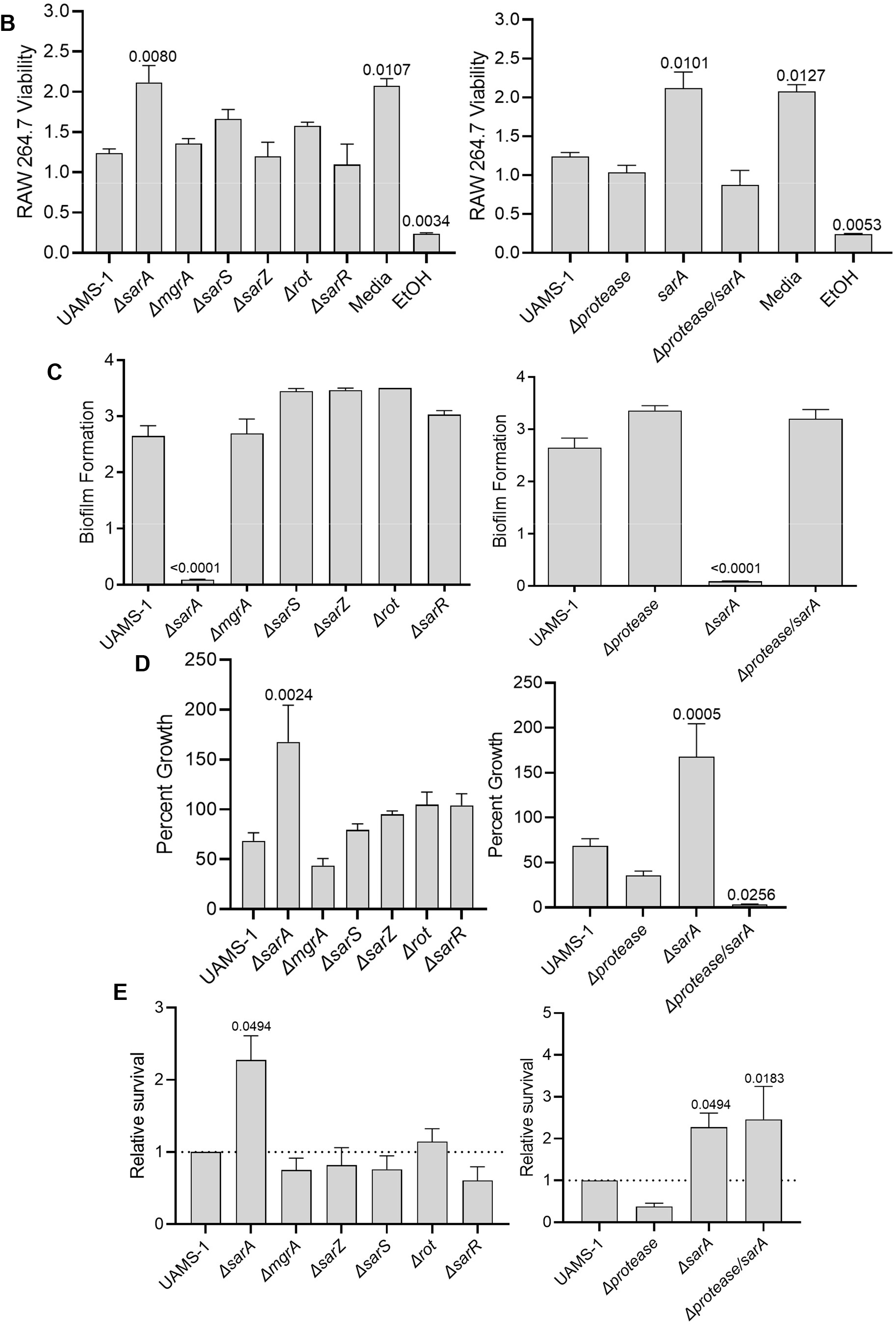
Mutation of *sarA* in UAMS-1 has the greatest effect on osteomyelitis related phenotypes owing primarily to its impact on protease production. A and B) Cytotoxicity of CM from overnight cultures of UAMS-1 and isogenic mutants was assessed using MC3T3-E1 (A) and RAW264.7 cells (B) as surrogates for osteoblasts and osteoclasts, respectively. CM, sterile bacterial culture media (negative control), or ethanol (positive control) was mixed in a 1:1 ratio with the appropriate cell culture medium for cytotoxicity assays. C) Biofilm formation was assessed using a microtiter plate assay. D) For each strain, 1×10^6^ cfu was inoculated into TSB with or without 10 μg/mL indolicidin and incubated overnight with shaking. Growth was measured by OD_600_ relative to a DMSO control for each mutant. E) 1×10^5^ cfu of bacterial cells from exponential phase cultures were mixed with 1.0 ml of whole human blood. A sample was taken immediately after mixing and after a 3 hr incubation. Percent survival of each mutant was calculated and standardized relative to the results observed with UAMS-1. The difference between the results observed with the *sarA* mutant and its protease-deficient derivative were not statistically significant. In all cases, statistical significance was assessed by one-way ANOVA. Numbers indicate p-values by comparison to the results observed with the UAMS-1 parent strain.

## Discussion

*Staphylococcus aureus* is arguably the most diverse of all bacterial pathogens given its ability to cause such a wide array of infections [43]. Among the many virulence factors which mediate this diversity are the extracellular proteases aureolysin, SspA/SspB, ScpA, or SplA-F, which have been shown to promote nutrient acquisition, tissue invasion, and evasion of host defenses [28,29,44–46]. However, recent reports have demonstrated that eliminating protease production results in increased virulence, while mutants that exhibit increased protease production also exhibit reduced virulence [22–25,38,47]. Neither of these would be predicted for a classic virulence factor.

This apparent paradox can be explained by the impact of these proteases on the virulence factor repertoire of *S. aureus*. Specifically, eliminating protease production increases virulence because it results in the increased abundance of other virulence factors [38,47]. Conversely, increased protease production has been shown to decrease virulence because it results in the decreased abundance of these virulence factors [21,25,26,30,48]. Such results suggest that extracellular proteases also serve a key post-translational regulatory role that must be kept in check to allow the proteases to serve their intended purposes on behalf of the bacterium without compromising the rest of the *S. aureus* virulence factor repertoire.

The importance of this post-translational control is reinforced by the number of regulatory loci that have been implicated in modulation of extracellular protease production [22,26,33,37,40]. However, it is impossible to put the relative role of these regulatory loci into context because few reports have included direct comparisons. It is also not possible from these reports to determine whether the impact of different regulatory loci occurs via a direct or indirect mechanism, particularly given the complexity and highly interactive nature of *S. aureus* regulatory circuits [49,50]. To begin to address these issues, we used the promoters associated with the genes and/or operons encoding aureolysin, ScpA, SspA/B, and SplA-F as DNA baits to capture proteins from whole cell lysates of S. *aureus*, which demonstrated that MgrA, Rot, SarA, SarR, SarS, and SarZ were captured by at least 3 of 4 protease-associated baits [23]. SarA was the most abundant protein captured by all four baits, and subsequent *in vitro* phenotypic comparisons of the corresponding mutants confirmed that mutation of *sarA* resulted in a greater increase in protease production that mutation of any of the other regulatory loci, irrespective of the functional status of the other loci [23].

These results suggest that *sarA* plays a major role in limiting the production of extracellular proteases and that, by doing so, plays a predominant role in mediating post-translational regulation in *S. aureus*. However, these studies were limited to *in vitro* experiments and therefore do not necessarily reflect *in vivo* significance, particularly in a unique microenvironment like the bone. Thus, our goal in these experiments was to assess the *in vivo* relevance of the genes encoding the regulatory proteins captured in our earlier experiments (*mgrA*, *rot*, *sarA*, *sarR*, *sarS*, and *sarZ*). We chose to use a murine osteomyelitis model based on our specific interest in overcoming the therapeutic recalcitrance of orthopaedic infections and because we had previously shown that increased protease production is correlated with decreased virulence in this model [22,25,48,51]. We also chose to include mutants generated in both the methicillin-resistant USA300 strain LAC and the methicillin-sensitive USA200 strain UAMS-1 (ATCC 49230). Isolates of the USA300 clonal lineage are clinically important as evidenced by the emergence of serious community-acquired infections [52–55]. However, they are also unique by comparison to other clinically relevant clonal lineages of *S. aureus* in several respects including gene content and gene expression [56,57]. UAMS-1 was isolated directly from the bone of an osteomyelitis patient undergoing surgical debridement and is demonstrably different from LAC at both a genotypic and phenotypic level [58,59]. The inclusion of such divergent clinical isolates increases the significance of the experiments we report in that it allowed us to determine whether we could identify common elements for potential therapeutic intervention.

In LAC, the results of our studies were very clear in that mutation of *sarA* resulted in a decrease in osteomyelitis virulence as reflected by cortical bone destruction, reactive new bone formation, bacterial burdens in the femur, and overall osteomyelitis scores, while none of the other regulatory mutants examined impacted any of these phenotypes to a statistically significant degree. This result could be correlated with cytotoxicity for osteoblasts and osteoclasts, which was reduced in CM from a LAC *sarA* mutant but was not altered with CM from any other regulatory mutant. Moreover, the reduced cytotoxicity of CM from the *sarA* mutant was a direct function of the increased production of extracellular proteases as evidenced by the fact cytotoxicity was restored by eliminating the ability of the *sarA* mutant to produce aureolysin, ScpA, SspA, SspB, and the spl-encoded proteases (SplA-F). Biofilm formation was also limited to a significant extent in the LAC *sarA* mutant and restored in the protease-deficient *sarA* mutant. Mutation of *rot* reduced biofilm formation in LAC in a protease-dependent manner, although to a lesser extent than mutating *sarA*.

Sensitivity to the antimicrobial peptide indolicidin was also decreased in the LAC *sarA* mutant in a protease-dependent manner. In contrast, sensitivity to indolicidin was increased in a LAC *sarZ* mutant. The reasons for this remain to be determined, but it has been shown that mutation of *sarZ* results in the increased transcription of *sarA* and decreased transcription of the gene encoding *sspA* [60]. Interestingly, survival in whole human blood was also increased to a statistically significant extent in the LAC *sarA* mutant, and this was not the case with any other regulatory mutant. However, eliminating protease production did not reverse this phenotype to a statistically significant extent, possibly suggesting that other loci regulated by *sarA* are involved. Overall, these studies are consistent with the conclusions that *S. aureus* must carefully limit the production of extracellular proteases and that SarA plays a direct and predominant role in this regard.

The results observed with UAMS-1 were consistent but less definitive than those observed with LAC. Specifically, mutation of *sarA* in UAMS-1 limited virulence in our osteomyelitis model, but not to the same extent observed with a LAC *sarA* mutant. Mutation of *sarA* had the same effect in both strains on cytotoxicity, biofilm formation, sensitivity to indolicidin, and survival in whole human blood, and, as with a LAC *sarA* mutant, eliminating protease production reversed all of these phenotypes except survival in blood. Thus, the results we report confirm that *sarA* plays an important role *in vivo* in limiting protease production, and that it does so in diverse clinical isolates. However, significant strain-dependent differences were observed with respect to other regulatory loci.

For example, mutation of *sarZ* had no impact on indolicidin sensitivity in UAMS-1. Moreover, mutation of the genes encoding two other members of the SarA family of proteins, namely MgrA and Rot, had no significant impact on any of the *in vitro* phenotypes we examined, but did result in reduced virulence in our osteomyelitis model to an equal or even greater extent than mutation of *sarA*. The fact that mutation of *rot* in LAC limited biofilm formation in a protease-dependent manner is consistent with an earlier report that focused solely on LAC [33], but it remains unclear why mutation of *rot* significantly limits virulence in our osteomyelitis model in UAMS-1 but not in LAC. These results nevertheless emphasize the importance of considering diverse strains of *S. aureus* before drawing conclusions about the impact of specific regulatory loci and their contribution to potentially important phenotypes.

We have previously shown that mutation of *rot* in UAMS-1 does result in a modest but statistically significant increase in protease production [23], and the results we present do not preclude the possibility that increased protease production contributes to the reduced virulence of a UAMS-1 *rot* mutant in our osteomyelitis model. At the same time, Rot is known to directly impact the production of a variety of secreted virulence factors and cell surface proteins [61–63]. Moreover, mutation of *rot* has also been shown to impact the transcription of other regulatory loci including *saeR* and *sarS*, while *agr, rbf, sarA, sarS, sarU*, and *sigB* have all been shown to impact either the functional activity or production of Rot [37,64,65]. It is particularly notable that *sarU* is one of the regulatory loci implicated in the *rot* regulatory circuit as it, along with the adjacent regulatory gene *sarT*, is present in LAC but not in UAMS-1 [58].

Similarly, Gimza *et al* [37] concluded that *mgrA* is one of the primary regulators of protease production. It affects at least 9 other regulatory loci (*agr, arIRS, atIR, sarR, sarS, sarV, sarX, sarZ, sigB*) and can be regulated by at least 7 others (*agr, arIRS, mntR, rex, sarV, sarZ*, and *xdrA*), which are in turn regulated by each other and by other regulatory loci such as *codY, nsaRS, rsrR* and *sarT* [37,66–71]. Moreover, mutation of *mgrA* is known to impact other potentially important phenotypes including toxin and nuclease production [72–74], cell clumping [66], immune evasion [72–75], capsule formation [72,73,76], and mammalian cell invasion [77]. Thus, the reason(s) mutation of *mgrA* and *rot* limit the virulence of UAMS-1 in osteomyelitis remain to be determined, but, irrespective of the mechanism(s) involved, these results do not contradict the conclusion that *sarA* is the only regulatory locus among those we examined that is consistently associated with reduced virulence in diverse clinical isolates of S. *aureus*. These results emphasize the importance of *in vivo* comparative studies to determine the relative impact of regulatory loci in the context of both infection and the diversity among clinical isolates of S. *aureus*. They also suggest that *sarA* is one of if not the most promising regulatory loci for the development of anti-virulence therapies targeting osteomyelitis owing to its role in limiting protease production as an important means of post-translational regulation of the *S. aureus* virulence factor repertoire.

In conclusion, the results we report emphasize the importance of *in vivo* comparative studies to determine the relative impact of regulatory loci in the context of infection and the diversity among clinical isolates of S. *aureus*. They also demonstrate that mutation of *sarA* attenuates the virulence of both LAC and UAMS-1 to a greater and more consistent extent than mutation of any of the other regulatory loci we examined. The results of our cytotoxicity and biofilm studies with both LAC and UAMS-1 are consistent with the hypothesis that this attenuation can be attributed to the increased production of extracellular proteases in *sarA* mutants, and in fact this has been proven *in vivo* in our osteomyelitis model using LAC [25]. Studies to confirm that this is also the case in a UAMS-1 *sarA* mutant and to determine the relative contribution of specific proteases are ongoing. Most importantly, our results point to *sarA* as the most promising protease regulatory locus for the development of anti-virulence therapies targeting osteomyelitis.

## Materials and Methods

### Bacterial strains and growth conditions

LAC and UAMS-1 regulatory mutants were generated by phage-mediated transduction using mutants available in the NTML as donor strains [23]. Derivatives of each parent strain and specific regulatory mutants with additional mutations in the genes encoding aureolysin, ScpA, SspA, SspB, and the Spl proteases were generated as previously described [24,30]. All strains were maintained as stocks at −80°C in tryptic soy broth (TSB) supplemented with 25% (v/v) glycerol. Bacteria were recovered from frozen stocks by plating on tryptic soy agar (TSA) containing appropriate antibiotics. Antibiotics were used at the following concentrations: chloramphenicol, 10 μg/ml; kanamycin, 50 μg/ml; neomycin, 50 μg/ml; erythromycin, 10 μg/ml; spectinomycin, 1 mg/ml; and tetracycline, 5 μg/ml. Prior to *in vivo* analysis, each strain was grown overnight at 37°C in TSB with shaking and without antibiotic selection, washed 3 times with sterile phosphate-buffered saline (PBS), and then resuspended in PBS at a density of 5 X 10^8^ colony-forming units (cfu) per ml. The concentration of each strain was confirmed by plating serial dilutions on TSA with and without appropriate antibiotic selection. 2 μl of this suspension (1 X 10^6^ cfu) was then used to infect mice.

### Murine osteomyelitis model

All experiments involving animals were reviewed and approved by the University of Arkansas for Medical Sciences Institutional Animal Care and Use Committee under animal use protocol number 4124. Experiments were performed as previously described and in accordance with NIH guidelines, the Animal Welfare Act, and United States federal law [25,51]. Briefly, 6-8 week-old C57BL/6 mice were anesthetized and the femur exposed by making an incision in the right hind limb. A unicortical defect was created in the middle of the exposed femur. 1 × 10^6^ bacterial cells prepared as described above were then injected into the medullary canal in a total volume of 2 μl. Muscle and skin were sutured, and the infection allowed to proceed for 14 days. Mice were humanely euthanized and the infected femurs recovered. After removing soft tissues, femurs were frozen at −80°C before imaging by microcomputed tomography (μCT). After imaging, femurs were homogenized and bacterial burdens determined as detailed below. At least two independent experiments with 5 mice per experimental group were done with LAC, UAMS-1, and their isogenic mutants.

### Microcomputed tomography (μCT)

In the first set of experiments done with each set of strains, image acquisition was done using a Skyscan 1174 X-ray Microtomograph (Bruker, Kontich, Belgium) with an isotropic voxel size of 6.7 μm, an X-ray voltage of 50 kV (800 μA) and a 0.25 mm aluminum filter [22,25]. Reconstruction was carried out using the Skyscan Nrecon software. The reconstructed cross-sectional slices were processed using the Skyscan CT-analyzer software as previously described to delineate regions of interest (ROIs) where reactive new bone (callus) was isolated from cortical bone [22,25]. The ROIs were used to calculate the volume of cortical bone, and the amount of cortical bone destruction was estimated by subtracting the value obtained from each bone from the average obtained from sham operated bones inoculated with PBS. New bone formation was quantified using the subtractive ROI function on the previously delineated cortical bone-including ROI images and calculating the bone volume included in the newly defined ROI.

In the second set of experiments, image acquisition was done using a Skyscan 1275 X-ray Microtomograph (Bruker, Kontich, Belgium) with an isotropic voxel size of 6.8 μm and an X-ray voltage of 40 kV (100 μA). Reconstruction was carried out using the Skyscan Nrecon software. The reconstructed cross-sectional slices were processed using the Skyscan CT-analyzer software to perform a semiautomated protocol to create preliminary ROIs of only cortical bone. The semi-automated protocol was as follows: global thresholding (low = 90; high = 255), round closing in 3D space pixel size 4, round opening in 3D space, pixel size 1, round closing in 3D space, pixel size 8, and round dilation in 3D space pixel size 3. The resulting images were loaded as ROI and corrected by drawing inclusive or exclusive contours on the periosteal surface to keep only the cortical bone. Using these defined ROIs, the volume of cortical bone was calculated using a threshold of 70-255, and the amount of cortical bone destruction estimated by subtracting the value obtained from each bone from the average obtained from sham operated bones inoculated with sterile PBS. New bone formation was quantified using the subtractive ROI function on the previously delineated cortical bone ROI images and calculating the bone volume included in the newly defined ROI using a threshold of 45-135.

In both sets of experiments, statistical analysis was done by one-way ANOVA with Dunnett’s correction. Comparisons were made with all mutants relative to the appropriate parent strain. A p-value ≤0.05 was considered statistically significant.

### Bacterial burdens in the femur

Immediately after μCT imaging, each femur was homogenized using a Bullet Blender 5 Gold (Next Advance Inc., Troy NY) and resuspended in 1 ml of PBS. Serial dilutions were then plated on TSA without antibiotic selection. In all experiments, the identity of the *S. aureus* strain isolated at the end of the experiment was confirmed as the same strain used to initiate the infection by polymerase chain reaction (PCR) analysis to confirm the presence of the *cna* gene in UAMS-1 and its absence in LAC, the presence of the *pvl* genes in LAC and their absence in UAMS-1, and the presence of the mutation under study. Colony forming units (cfu) were counted and differences between groups assessed using a one-way analysis of variance (ANOVA) model. Briefly, cfu data was logarithmically transformed prior to analysis. For samples with no bacterial counts, 1 cfu was used so the samples would be included in the logarithmically transformed datasets. Contrasts were defined to assess the comparisons of interest. Adjustments for multiple comparisons were made using simultaneous general linear hypothesis testing procedures. Analyses were done using R (version 3.4.3, R Foundation for Statistical Computing, Vienna, Austria). Multiple comparison procedures were implemented using the R library multicomp. Adjusted p-values ≤0.05 were considered significant.

### Osteomyelitis score

In some cases, quantitative μCT analysis was not possible because the femur was broken. We addressed this based on the premise that fracture was indicative of pathology by developing an osteomyelitis (OM) score for each experimental animal. Therefore, in the absence of fracture, the score formula was based on the sum of the amount of cortical bone destruction (mm^3^) + the amount of reactive bone formation (mm^3^) + the log_10_ of the cfu per femur. In those mice in which the bone was fractured, the numbers used for cortical bone destruction and reactive bone formation were derived by adding one standard deviation to the highest scores observed with an intact bone from the same experimental group. Statistical analysis was done by comparing OM scores for individual mice in each experimental group by one-way ANOVA with Dunnett’s correction. A p-value ≤0.05 was considered statistically significant.

### Cytotoxicity for mammalian cells

RAW 264.7 and MC3T3-E1 cells were used as surrogates for osteoclasts and osteoblasts, respectively. Cytotoxicity was assessed using sterile conditioned medium (CM) from overnight bacterial cultures as previously described [22,23]. Briefly, RAW264.7 and MC3T3-E1 cell lines were obtained from the American Type Culture Collection (ATCC). RAW264.7 cells were grown in Dulbecco’s Modified Eagle’s Medium (D-MEM), while MC3T3-E1 cells were grown in Alpha Minimum Essential Medium (αMEM). Culture media was supplemented with 10% fetal bovine serum and penicillin and streptomycin (100 μg/ml each). Cells were grown at 37°C in 5% CO2 with the replacement of the media as needed every 2 to 3 days. For cytotoxicity assays, cells were seeded into black clear-bottom 96-well tissue culture-grade plates at a density of 10,000 cells per well for MC3T3-E1 cells or 50,000 cells per well for RAW 264.7 cells. After 24 h, the growth medium was removed and replaced with medium containing a 1:1 mixture of cell culture medium with antibiotics and *S. aureus* CM. Monolayers were incubated for an additional 24 h prior to removal of the medium and assessment of cell viability using the calcein-AM LIVE/DEAD Viability/Cytotoxicity Kit (Thermo Fisher Scientific) according to the manufacturer’s specifications. Fluorescence intensity was read on a plate reader with an excitation wavelength of 485 nm and emission wavelength of 520 nm with the gain set to 95% of the intensity observed in the well that exhibited the highest fluorescence intensity. Cell viability is reported as the fluorescence intensity divided by 100,000. A 1:1 mixture of cell culture medium and TSB or ethanol (EtOH) was used as negative and positive cytotoxicity controls, respectively.

### Sensitivity to indolicidin

Sensitivity to the antimicrobial peptide indolicidin was assessed as previously described [47,78] with modification. Specifically, each strain was grown overnight in TSB at 37°C with shaking. Cultures were standardized in TSB and 1 x 10^6^ cfu added to the wells of a 96-well microtiter plate containing 50 μl of 2-fold concentrated TSB, 30 μl of sterile water, and 20 μl of 50 μg/ml indolicidin in DMSO. DMSO without indolicidin was used as a control. After overnight incubation at 37°C with shaking, optical density at 600 nm (OD_600_) was determined using a microtiter plate reader and the percent growth calculated as the OD_600_ in the presence of indolicidin divided by the OD_600_ without indolicidin X 100. Results are reported as the average of 3 biological replicates, each of which included 3 experimental replicates.

### Survival in whole human blood

Survival in whole human blood was performed as previously described [47,78] with minor modification. Briefly, strains were grown overnight in TSB at 37°C with shaking, diluted 1:100 in 4 ml TSB, and incubated for an additional 3 hours. Cells from 1 ml of each culture were harvested by centrifugation, washed twice in PBS, and standardized to an equivalent optical density in PBS. 1 x 10^6^ cfu was then added to 1 ml of whole human blood (BioIVT). An aliquot was immediately removed, serially diluted, and plated to verify the inoculum. The mixture was then incubated at 37° with shaking for 3 h, after which another aliquot was removed, diluted, and plated. Relative percent survival was determined by finding the percent survival of each mutant [(cfu at 3 hours divided by cfu at 0 hours] X 100) and dividing by the percent survival of the appropriate parent strain in each experiment. Experiments were done as at least 3 biological replicates, each with 3 experimental replicates.

## Acknowledgments

We would like to acknowledge Horace “Trey” J. Spencer in the University of Arkansas for Medical Sciences Department of Biostatistics for his assistance with the statistical analysis and the Center for Microbial Pathogenesis and Host Inflammatory Responses Cellular Imaging and Molecular Biology Cores at the University of Arkansas for Medical Sciences.

## References

1. Hatzenbeuhler J, Pulling TJ. Diagnosis and Management of Osteomyelitis. Am Fam Physician. 2011;84: 1027–1033. Available: https://www.aafp.org/pubs/afp/issues/2011/1101/p1027.html

2. Nasser A, Azimi T, Ostadmohammadi S, Ostadmohammadi S. A comprehensive review of bacterial osteomyelitis with emphasis on Staphylococcus aureus. Microb Pathog. 2020;148: 104431. doi:10.1016/J.MICPATH.2020.104431

3. Veis DJ, Cassat JE. Infectious Osteomyelitis: Marrying Bone Biology and Microbiology to Shed New Light on a Persistent Clinical Challenge. 2021. doi:10.1002/jbmr.4279

4. Lew DP, Waldvogel FA. Osteomyelitis. Lancet. 2004;364: 369–379. doi:10.1016/S0140-6736(04)16727-5

5. Urish KL, Cassat JE. Staphylococcus aureus Osteomyelitis: Bone, Bugs, and Surgery. Infect Immun. 2021;88: e00932–19. doi:10.1128/IAI.00932-19

6. Cierny G. Surgical Treatment of Osteomyelitis. Plast Reconstr Surg. 2011; 127: 190S–204S. doi:10.1097/PRS.0b013e3182025070

7. Brady RA, Leid JG, Calhoun JH, Costerton JW, Shirtliff ME. Osteomyelitis and the role of biofilms in chronic infection. FEMS Immunol Med Microbiol. 2008;52: 13–22. doi:10.1111/J.1574-695X.2007.00357.X

8. Masters EA, Trombetta RP, de Mesy Bentley KL, Boyce BF, Gill AL, Gill SR, et al. Evolving concepts in bone infection: redefining “biofilm”, “acute vs. chronic osteomyelitis”, “the immune proteome” and “local antibiotic therapy.” Bone Res. 2019;7: 20. doi:10.1038/s41413-019-0061-z

9. Zimmerli W, Sendi P. Orthopaedic biofilm infections. APMIS. 2017;125: 353–364. doi:10.1111/APM.12687

10. Bongiorno D, Musso N, Lazzaro LM, Mongelli G, Stefani S, Campanile F. Detection of methicillin-resistant Staphylococcus aureus persistence in osteoblasts using imaging flow cytometry. Microbiologyopen. 2020;9: e1017. doi:10.1002/MBO3.1017

11. Gimza BD, Cassat JE. Mechanisms of Antibiotic Failure During Staphylococcus aureus Osteomyelitis. Front Immunol. 2021;12: 638085. doi:10.3389/fimmu.2021.638085

12. Krauss JL, Roper PM, Ballard A, Shih C-C, Fitzpatrick JAJ, Cassat JE, et al. Staphylococcus aureus Infects Osteoclasts and Replicates Intracellularly. mBio. 2019;10. doi:10.1128/MBIO.02447-19

13. Mouton W, Josse J, Jacqueline C, Abad L, Trouillet-Assant S, Caillon J, et al. Staphylococcus aureus internalization impairs osteoblastic activity and early differentiation process. Scientific Reports 2021 11:1. 2021;11: 1–10. doi:10.1038/s41598-021-97246-y

14. Ford CA, Cassat JE. Measurement of Osteoblast Cytotoxicity Induced by S. aureus Secreted Toxins. Methods Mol Biol. 2021;2341: 141–152. doi:10.1007/978-1-0716-1550-8_17

15. Rao N, Ziran BH, Lipsky BA. Treating osteomyelitis: Antibiotics and surgery. Plast Reconstr Surg. 2011;127. doi:10.1097/PRS.0B013E3182001F0F

16. del Pozo JL, Patel R. The Challenge of Treating Biofilm-associated Bacterial Infections. Clin Pharmacol Ther. 2007;82: 204–209. doi:10.1038/SJ.CLPT.6100247

17. Atwood DN, Beenken KE, Lantz TL, Meeker DG, Lynn WB, Mills WB, et al. Regulatory Mutations Impacting Antibiotic Susceptibility in an Established Staphylococcus aureus Biofilm. Antimicrob Agents Chemother. 2016;60: 1826. doi:10.1128/AAC.02750-15

18. Beenken KE, Blevins JS, Smeltzer MS. Mutation of sarA in Staphylococcus aureus Limits Biofilm Formation. Infect Immun. 2003;71: 4206–4211. doi:10.1128/IAI.71.7.4206-4211.2003

19. Weiss EC, Spencer HJ, Daily SJ, Weiss BD, Smeltzer MS. Impact of sarA on antibiotic susceptibility of Staphylococcus aureus in a catheter-associated in vitro model of biofilm formation. Antimicrob Agents Chemother. 2009;53: 2475–2482. doi:10.1128/AAC.01432-08/ASSET/C80A6294-C54D-4CB6-BCAD-6CB1B331160C/ASSETS/GRAPHIC/ZAC0060981210008.JPEG

20. Weiss EC, Zielinska A, Beenken KE, Spencer HJ, Daily SJ, Smeltzer MS. Impact of sarA on daptomycin susceptibility of Staphylococcus aureus biofilms in vivo. Antimicrob Agents Chemother. 2009;53: 4096–4102. doi:10.1128/AAC.00484-09/ASSET/E48BE7BA-1456-48F3-90FD-7F5520DF409B/ASSETS/GRAPHIC/ZAC0100984890007.JPEG

21. Alkam D, Jenjaroenpun P, Ramirez AM, Beenken KE, Spencer HJ, Smeltzer MS. The Increased Accumulation of Staphylococcus aureus Virulence Factors Is Maximized in a purR Mutant by the Increased Production of SarA and Decreased Production of Extracellular Proteases. Infect Immun. 2021;89. doi:10.1128/IAI.00718-20

22. Loughran AJ, Gaddy D, Beenken KE, Meeker DG, Morello R, Zhao H, et al. Impact of sarA and phenol-soluble modulins on the pathogenesis of osteomyelitis in diverse clinical isolates of Staphylococcus aureus. Infect Immun. 2016;84: 2586–2594. doi:10.1128/IAI.00152-16/SUPPL_FILE/ZII999091802SD1.XLSX

23. Ramirez AM, Beenken KE, Byrum SD, Tackett AJ, Shaw LN, Gimza BD, et al. SarA plays a predominant role in controlling the production of extracellular proteases in the diverse clinical isolates of Staphylococcus aureus LAC and UAMS-1. Virulence. 2020;11: 1738–1762. doi:10.1080/21505594.2020.1855923

24. Rom JS, Beenken KE, Ramirez AM, Walker CM, Echols EJ, Smeltzer MS. Limiting protease production plays a key role in the pathogenesis of the divergent clinical isolates of Staphylococcus aureus LAC and UAMS-1. Virulence. 2021;12: 584–600. doi:10.1080/21505594.2021.1879550

25. Ramirez AM, Byrum SD, Beenken KE, Washam C, Edmondson RD, Mackintosh SG, et al. Exploiting Correlations between Protein Abundance and the Functional Status of saeRS and sarA To Identify Virulence Factors of Potential Importance in the Pathogenesis of Staphylococcus aureus Osteomyelitis. ACS Infect Dis. 2020;6: 237–249. doi:10.1021/acsinfecdis.9b00291

26. Byrum SD, Loughran AJ, Beenken KE, Orr LM, Storey AJ, Mackintosh SG, et al. Label-Free Proteomic Approach to Characterize Protease-Dependent and-Independent Effects of sarA Inactivation on the Staphylococcus aureus Exoproteome. J Proteome Res. 2018;17: 3384–3395. doi:10.1021/acs.jproteome.8b00288

27. Tsang LH, Cassat JE, Shaw LN, Beenken KE, Smeltzer MS. Factors Contributing to the Biofilm-Deficient Phenotype of Staphylococcus aureus sarA Mutants. PLoS One. 2008;3: e3361. doi:10.1371/JOURNAL.PONE.0003361

28. Lehman MK, Nuxoll AS, Yamada KJ, Kielian T, Carson SD, Feya PD. Protease-Mediated Growth of Staphylococcus aureus on Host Proteins Is opp3 Dependent. mBio. 2019;10. doi:10.1128/MBIO.02553-18

29. Tam K, Torres VJ. Staphylococcus aureus Secreted Toxins and Extracellular Enzymes. Microbiol Spectr. 2019;7: 10.1128/microbiolspec.GPP3-0039-2018. doi:10.1128/microbiolspec.GPP3-0039-2018

30. Zielinska AK, Beenken KE, Mrak LN, Spencer HJ, Post GR, Skinner RA, et al. sarA-mediated repression of protease production plays a key role in the pathogenesis of Staphylococcus aureus USA300 isolates. Mol Microbiol. 2012;86: 1183–1196. doi:10.1111/mmi.12048

31. Mootz JM, Malone CL, Shaw LN, Horswill AR. Staphopains Modulate Staphylococcus aureus Biofilm Integrity. 2013 [cited 17 Nov 2021]. doi:10.1128/IAI.00377-13

32. Rom JS, Atwood DN, Beenken KE, Meeker DG, Loughran AJ, Spencer HJ, et al. Impact of Staphylococcus aureus regulatory mutations that modulate biofilm formation in the USA300 strain LAC on virulence in a murine bacteremia model. Virulence. 2017/10/04. 2017;8: 1776–1790. doi:10.1080/21505594.2017.1373926

33. Mootz JM, Benson MA, Heim CE, Crosby HA, Kavanaugh JS, Dunman PM, et al. Rot is a key regulator of Staphylococcus aureus biofilm formation. Mol Microbiol. 2015;96: 388–404. doi:10.1111/MMI.12943/SUPPINFO

34. Park JH, Lee JH, Cho MH, Herzberg M, Lee J. Acceleration of protease effect on Staphylococcus aureus biofilm dispersal. FEMS Microbiol Lett. 2012;335: 31–38. doi:10.1111/J.1574-6968.2012.02635.X

35. Sonesson A, Przybyszewska K, Eriksson S, Mörgelin M, Kjellström S, Davies J, et al. Identification of bacterial biofilm and the Staphylococcus aureus derived protease, staphopain, on the skin surface of patients with atopic dermatitis. Sci Rep. 2017;7. doi:10.1038/S41598-017-08046-2

36. Rom JS, Ramirez AM, Beenken KE, Sahukhal GS, Elasri MO, Smeltzer MS. The Impacts of msaABCR on sarA-Associated Phenotypes Are Different in Divergent Clinical Isolates of Staphylococcus aureus. Infect Immun. 2020;88. doi:10.1128/IAI.00530-19

37. Gimza BD, Larias MI, Budny BG, Shaw LN. Mapping the Global Network of Extracellular Protease Regulation in Staphylococcus aureus. mSphere. 2019;4: e00676–19. doi:10.1128/mSphere.00676-19

38. Gimza BD, Jackson JK, Frey AM, Budny BG, Chaput D, Rizzo DN, et al. Unraveling the Impact of Secreted Proteases on Hypervirulence in Staphylococcus aureus. mBio. 2021;12: 1–15. doi:10.1128/MBIO.03288-20

39. Bose JL, Fey PD, Bayles KW. Genetic tools to enhance the study of gene function and regulation in Staphylococcus aureus. Appl Environ Microbiol. 2013/01/25. 2013;79: 2218–2224. doi:10.1128/AEM.00136-13

40. Fey PD, Endres JL, Yajjala VK, Widhelm TJ, Boissy RJ, Bose JL, et al. A Genetic Resource for Rapid and Comprehensive Phenotype Screening of Nonessential Staphylococcus aureus Genes. mBio. 2013;4. doi:10.1128/MBIO.00537-12

41. Balasubramanian D, Harper L, Shopsin B, Torres VJ. Staphylococcus aureus pathogenesis in diverse host environments. Pathog Dis. 2017;75. doi:10.1093/FEMSPD/FTX005

42. Wilde AD, Snyder DJ, Putnam NE, Valentino MD, Hammer ND, Lonergan ZR, et al. Bacterial Hypoxic Responses Revealed as Critical Determinants of the Host-Pathogen Outcome by TnSeq Analysis of Staphylococcus aureus Invasive Infection. PLoS Pathog. 2015;11: e1005341. doi:10.1371/JOURNAL.PPAT.1005341

43. Lowy FD. Staphylococcus aureus infections. N Engl J Med. 1998;339: 520–532. doi:10.1056/NEJM199808203390806

44. Kantyka T, Pyrc K, Gruca M, Smagur J, Plaza K, Guzik K, et al. Staphylococcus aureus Proteases Degrade Lung Surfactant Protein A Potentially Impairing Innate Immunity of the Lung. J Innate Immun. 2013;5: 251. doi:10.1159/000345417

45. Pietrocola G, Nobile G, Rindi S, Speziale P. Staphylococcus aureus Manipulates Innate Immunity through Own and Host-Expressed Proteases. Front Cell Infect Microbiol. 2017;7: 166. doi:10.3389/FCIMB.2017.00166

46. Singh V, Phukan UJ. Interaction of host and Staphylococcus aureus protease-system regulates virulence and pathogenicity. Medical Microbiology and Immunology 2018 208:5. 2018;208: 585–607. doi:10.1007/S00430-018-0573-Y

47. Kolar SL, Ibarra JA, Rivera FE, Mootz JM, Davenport JE, Stevens SM, et al. Extracellular proteases are key mediators of Staphylococcus aureus virulence via the global modulation of virulence-determinant stability. Microbiologyopen. 2013;2: 18–34. doi:10.1002/mbo3.55

48. Cassat JE, Hammer ND, Campbell JP, Benson MA, Perrien DS, Mrak LN, et al. A secreted bacterial protease tailors the Staphylococcus aureus virulence repertoire to modulate bone remodeling during osteomyelitis. Cell Host Microbe. 2013;13: 759–772. doi:10.1016/j.chom.2013.05.003

49. Jenul C, Horswill AR. Regulation of Staphylococcus aureus virulence. Microbiol Spectr. 2018;6. doi:10.1128/MICROBIOLSPEC.GPP3-0031-2018

50. Priest NK, Rudkin JK, Feil EJ, van den Elsen JMH, Cheung A, Peacock SJ, et al. From genotype to phenotype: can systems biology be used to predict Staphylococcus aureus virulence? Nat Rev Microbiol. 2012;10: 791. doi:10.1038/NRMICRO2880

51. Beenken KE, Mrak LN, Zielinska AK, Atwood DN, Loughran AJ, Griffin LM, et al. Impact of the functional status of saeRS on in vivo phenotypes of Staphylococcus aureus sarA mutants. Mol Microbiol. 2014/05/12. 2014;92: 1299–1312. doi:10.1111/mmi.12629

52. Hidron AI, Low CE, Honig EG, Blumberg HM. Emergence of community-acquired meticillin-resistant Staphylococcus aureus strain USA300 as a cause of necrotising community-onset pneumonia. Lancet Infect Dis. 2009;9: 384–392. doi:10.1016/S1473-3099(09)70133-1

53. Mediavilla JR, Chen L, Mathema B, Kreiswirth BN. Global epidemiology of community-associated methicillin resistant Staphylococcus aureus (CA-MRSA). Curr Opin Microbiol. 2012;15: 588–595. doi:10.1016/J.MIB.2012.08.003

54. Nimmo GR. USA300 abroad: global spread of a virulent strain of community-associated methicillin-resistant Staphylococcus aureus. Clinical Microbiology and Infection. 2012;18: 725–734. doi:10.1111/J.1469-0691.2012.03822.X

55. Thurlow LR, Joshi GS, Richardson AR. Virulence strategies of the dominant USA300 lineage of community-associated methicillin-resistant Staphylococcus aureus (CA-MRSA). FEMS Immunol Med Microbiol. 2012;65: 5–22. doi:10.1111/J.1574-695X.2012.00937.X

56. Diep BA, Gill SR, Chang RF, Phan TH, Chen JH, Davidson MG, et al. Complete genome sequence of USA300, an epidemic clone of community-acquired meticillin-resistant Staphylococcus aureus. Lancet. 2006;367: 731–739. doi:10.1016/S0140-6736(06)68231-7

57. Otto M. Staphylococcus aureus toxins. Curr Opin Microbiol. 2014;17: 32–37. doi:10.1016/j.mib.2013.11.004

58. Cassat JE, Dunman PM, McAleese F, Murphy E, Projan SJ, Smeltzer MS. Comparative genomics of Staphylococcus aureus musculoskeletal isolates. J Bacteriol. 2005;187: 576–592. doi:10.1128/JB.187.2.576-592.2005

59. Gillaspy AF, Hickmon SG, Skinner RA, Thomas JR, Nelson CL, Smeltzer MS. Role of the accessory gene regulator (agr) in pathogenesis of staphylococcal osteomyelitis. Infect Immun. 1995;63: 3373–3380. doi:10.1128/iai.63.9.3373-3380.1995

60. Tamber S, Cheung AL. SarZ Promotes the Expression of Virulence Factors and Represses Biofilm Formation by Modulating SarA and agr in Staphylococcus aureus. Infect Immun. 2009;77: 419. doi:10.1128/IAI.00859-08

61. Saïd-Salim B, Dunman PM, McAleese FM, Macapagal D, Murphy E, McNamara PJ, et al. Global Regulation of Staphylococcus aureus Genes by Rot. J Bacteriol. 2003;185: 610. doi:10.1128/JB.185.2.610-619.2003

62. Mcnamara PJ, Milligan-Monroe KC, Khalili S, Proctor RA. Identification, cloning, and initial characterization of rot, a locus encoding a regulator of virulence factor expression in Staphylococcus aureus. J Bacteriol. 2000;182: 3197–3203. doi:10.1128/JB.182.11.3197-3203.2000/ASSET/21DA62A8-DDD9-448D-9A17-4D14EE2D50E2/ASSETS/GRAPHIC/JB1101770005.JPEG

63. Benson MA, Lilo S, Wasserman GA, Thoendel M, Smith A, Horswill AR, et al. Staphylococcus aureus regulates the expression and production of the staphylococcal superantigen-like secreted proteins in a Rot-dependent manner. Mol Microbiol. 2011;81: 659–675. doi:10.1111/J.1365-2958.2011.07720.X

64. Oscarsson J, Tegmark-Wisell K, Arvidson S. Coordinated and differential control of aureolysin (aur) and serine protease (sspA) transcription in Staphylococcus aureus by sarA, rot and agr (RNAIII). International Journal of Medical Microbiology. 2006;296: 365–380. doi:10.1016/J.IJMM.2006.02.019

65. Hsieh HY, Ching WT, Stewart GC. Regulation of Rot Expression in Staphylococcus aureus. J Bacteriol. 2008;190: 546. doi:10.1128/JB.00536-07

66. Crosby HA, Schlievert PM, Merriman JA, King JM, Salgado-Pabón W, Horswill AR. The Staphylococcus aureus Global Regulator MgrA Modulates Clumping and Virulence by Controlling Surface Protein Expression. PLoS Pathog. 2016;12. doi:10.1371/JOURNAL.PPAT.1005604

67. Ingavale S, van Wamel W, Luong TT, Lee CY, Cheung AL. Rat/MgrA, a Regulator of Autolysis, Is a Regulator of Virulence Genes in Staphylococcus aureus. Infect Immun. 2005;73: 1423. doi:10.1128/IAI.73.3.1423-1431.2005

68. Manna AC, Ingavale SS, Maloney MB, van Wamel W, Cheung AL. Identification of sarV (SA2062), a New Transcriptional Regulator, Is Repressed by SarA and MgrA (SA0641) and Involved in the Regulation of Autolysis in Staphylococcus aureus. J Bacteriol. 2004;186: 5267. doi:10.1128/JB.186.16.5267-5280.2004

69. Manna AC, Cheung AL. Transcriptional regulation of the agr locus and the identification of DNA binding residues of the global regulatory protein SarR in Staphylococcus aureus. Mol Microbiol. 2006;60: 1289–1301. doi:10.1111/J.1365-2958.2006.05171.X

70. Manna AC, Ballal A, Ray B. sarZ, a sarA family gene, is transcriptionally activated by MgrA and is involved in the regulation of genes encoding exoproteins in Staphylococcus aureus. J Bacteriol. 2009;191: 1656–1665. doi:10.1128/JB.01555-08

71. Trotonda MP, Tamber S, Memmi G, Cheung AL. MgrA represses biofilm formation in Staphylococcus aureus. Infect Immun. 2008;76: 5645–5654. doi:10.1128/IAI.00735-08

72. Luong TT, Dunman PM, Murphy E, Projan SJ, Lee CY. Transcription Profiling of the mgrA Regulon in Staphylococcus aureus. J Bacteriol. 2006;188: 1899. doi:10.1128/JB.188.5.1899-1910.2006

73. Luong TT, Newell SW, Lee CY. Mgr, a novel global regulator in Staphylococcus aureus. J Bacteriol. 2003;185: 3703–3710. doi:10.1128/JB.185.13.3703-3710.2003

74. Li L, Wang G, Cheung A, Abdelhady W, Seidl K, Xiong YQ. MgrA Governs Adherence, Host Cell Interaction, and Virulence in a Murine Model of Bacteremia Due to Staphylococcus aureus. J Infect Dis. 2019;220: 1019. doi:10.1093/INFDIS/JIZ219

75. Jakub Kwiecinski AM, Kratofil RM, Parlet CP, Surewaard BG, Kubes P, Horswill Correspondence AR, et al. Staphylococcus aureus uses the ArlRS and MgrA cascade to regulate immune evasion during skin infection. Cell Rep. 2021;36. doi:10.1016/j.celrep.2021.109462

76. Gupta RK, Alba J, Xiong YQ, Bayer AS, Lee CY. MgrA activates expression of capsule genes, but not the α-toxin gene in experimental Staphylococcus aureus endocarditis. J Infect Dis. 2013;208: 1841–1848. doi:10.1093/INFDIS/JIT367

77. Lei MG, Lee CY. MgrA Activates Staphylococcal Capsule via SigA-Dependent Promoter. J Bacteriol. 2021;203. doi:10.1128/JB.00495-20

78. Kolar SL, Nagarajan V, Oszmiana A, Rivera FE, Miller HK, Davenport JE, et al. NsaRS is a cell-envelope-stress-sensing two-component system of Staphylococcus aureus. Microbiology (N Y). 2011;157: 2206. doi:10.1099/MIC.0.049692-0

